# Analysis of intracellular fatty acid metabolism during Doxorubicin induced senescence of MCF7 cells using Raman Imaging

**DOI:** 10.1101/2025.09.17.676702

**Authors:** Swarang Sachin Pundlik, Ashwin Venkateshvaran, Yavanica Suresh, Hitesh Mamgain, Arvind Ramanathan

## Abstract

Cellular senescence, a stable growth-arrested state induced by stress or chemotherapeutic agents, is accompanied by metabolic remodeling that supports the senescence-associated secretory phenotype (SASP). Among these pathways, lipid and arachidonic acid (AA) metabolism play central roles in maintaining and propagating the senescent state. Here, we used hyperspectral confocal Raman microscopy to visualize biochemical remodeling in MCF7 human breast adenocarcinoma cells undergoing doxorubicin-induced senescence. Raman spectral analysis and principal-component decomposition revealed time-dependent alterations in lipid-associated vibrational modes—particularly CH₂ and C=C stretching—consistent with enhanced lipid accumulation and remodeling between days 10 and 15 after DNA-damage induction. PCA of lipid-rich compartments isolated using true component analysis also confirms progressive increases in triacylglycerol and unsaturated lipid signatures. Using deuterated arachidonic acid (AA-d₈) and COX-2 inhibition, we further demonstrated real-time intracellular AA metabolism by tracking C=C–D stretching peaks (2220–2254 cm⁻¹) in the Raman-silent window. The ratio of these deuterium bands to CH₂ stretching provided a label-free quantitative metric for COX2–dependent AA turnover in senescent cells. Together, these findings establish Raman hyperspectral imaging as a powerful, non-perturbative tool to map lipid and oxylipin metabolism during cellular senescence, offering new avenues to identify metabolic vulnerabilities in senescent tumor cells.

## Introduction

Cellular senescence is a stable proliferative arrest induced by diverse stresses and characterized by a Senescence-Associated Secretory Phenotype (SASP). The SASP includes inflammatory cytokines, proteases, and growth factors, which influence tissue remodeling, aging, and tumorigenesis^1–3^. Cancer cells, including MCF7 human breast adenocarcinoma cells, undergo senescence upon treatment with chemotherapeutic compounds like Doxorubicin (Doxo)^4^. These senescent tumor cells alter the physiology of the surrounding cancerous and non-cancerous cells via paracrine and juxtacrine signaling mediated by the components of the SASP, leading to secondary tumorigenesis and other pathologies^5–7^.

Metabolic reprogramming is increasingly recognized as integral to senescence phenotypes, not merely a bystander^8^. Lipids lie at the heart of that metabolic shift. In replicative and stress-induced senescence models, lipidomic analysis showed accumulation of triacylglycerols enriched in polyunsaturated fatty acids and modulation of membrane lipid remodeling^9^. Transcriptome and lipidome integrative analyses point to upregulated lipid uptake and storage genes in senescent cells, including the fatty acid translocator CD36. In fact, forced expression of CD36 in proliferating cells induces SASP factors^10^. These data argue for a causal axis from lipid uptake/storage to SASP execution.

Arachidonic acid (AA) metabolism is a specialized lipid axis in senescent cell biology. Wiley et al. demonstrated that senescent cells activate oxylipin biosynthesis, including prostaglandins, which reinforce cell cycle arrest and SASP expression; they identified a specific product, dihomo- 15d-PGJ₂, as a senolysis biomarker^11^. Parallel work implicates COX-2 / PGE₂ signaling in driving senescence onset and maintenance, and leukotriene outputs from lung fibroblasts in fibrotic niche remodeling ^12,13^. Our previous study has shown that 15d-PGJ2, a metabolite of Arachidonic Acid secreted by senescent myoblasts upon DNA damage, inhibits the differentiation of skeletal myoblasts by altering intracellular HRas dynamics^14^. Collectively, these studies place AA-pathway lipids as both sensors and effectors of senescence.

To characterize the senescent state, investigators routinely use multiplexed markers. Classic hallmarks include activity of senescence-associated β-galactosidase (SA-β-gal) at pH 6^15^, upregulation of cyclin inhibitors p21^CIP1/WAF1^ and p16^INK4a^ ^16^, and loss or reduction of Lamin B1^17^. Persistent DNA damage foci (γH2AX) mark unresolved damage^18^. Additional markers include chromatin reorganization (SAHF), HMGB1 relocalization, mitochondrial dysfunction, and metabolic shifts^18,19^. Because single markers are often insufficiently specific, field consensus recommends using panels of markers.

Recent advances in vibrational microscopy now allow label-free imaging of biomolecular composition with subcellular resolution. Confocal Raman or coherent Raman modalities (such as CARS/SRS) can spatially resolve lipid droplets, membrane changes, nucleolar protein conformation, and other chemical microdomains without exogenous labels. Raman fingerprints are promising markers for studying senescence^20^. For senescent cells, coherent Raman imaging has been used to visualize nucleolar amide-I shifts correlated with β-sheet aggregation in binucleated senescent cells^21^. In this manuscript, we apply confocal Raman imaging to senescent cells in conjunction with multiplexed molecular marker validation (SA-β-gal, p21 expression, nuclear enlargement, flattening of the cell morphology). Our aims are: (i) to spatially localize lipid species and AA-derived metabolites in single senescent cells; (ii) to correlate Raman spectral features with canonical senescence markers; (iii) to propose label-free spectral metrics that reflect senescence burden or responses to senolysis. By anchoring imaging readouts to validated lipids and markers, we hope to establish a robust, non-perturbative phenotyping tool for senescent cell states that can integrate into metabolic and pharmacologic studies.

## Materials and Methods

### Cell Culture

- MCF7 Human Breast Adenocarcinoma cells (HTB-22) were maintained in DMEM (Gibco) supplemented with 10% Fetal Bovine Serum (FBS) (Gibco) and 1% Penicillin-Streptomycin (Gibco) (DMEM complete medium). @37°C, 5% CO2.
- Cells were trypsinized with 0.125% trypsin-EDTA solution (Gibco) and were seeded in glass-bottom dishes for experiments.

### Treatments

- **Doxorubicin**:
- `o` Doxorubicin (Sigma Aldrich) was dissolved in DMSO (10 mM). Doxo (10 mM) was diluted in DMSO to make an intermediate stock (100 μM).

`o` MCF7 cells were treated with Doxo (150 nM) by diluting Doxo in DMEM complete medium for 3 days.
`o` The media was changed on Day 3 with fresh DMEM complete medium containing Doxo.
`o` From Day 5, the cells were incubated in DMEM complete medium without Doxo till the end of the experiment (Day 10, 15, 18, and 25).
`o` MCF7 cells were seeded in glass-bottom dishes and incubated in DMEM complete medium without Doxo for 24 hours, and were used as the proliferating control.

### • AA-d8 and Cox2 inhibition

`o` 5,6,8,9,11,12,14,15 D8 – Arachidonic Acid (AA-d8) obtained from Cayman Chemical Company was reconstituted in DMSO (25 mM).
`o` Cay-10404, a Cox2 inhibitor, was obtained from Cayman Chemical Company and was dissolved in DMSO (30 mM).
`o` Senescent MCF7 cells after 15 days of treatment with Doxo were pre-treated with Cay-10404 (30 μM) in DMEM complete medium for 2 hours for the inhibition of Cox2 enzyme.
`o` The cells were then treated with Cay-10404 (30 μM) and AA-d8 (100 μM) in

DMEM complete medium overnight.

- The cells were washed with 1x Dulbecco’s Phosphate Buffer Saline (DPBS) (Gibco) and were treated with AA-d8 (100 μM) in DMEM complete medium without Cay-10404 to allow resumption of Cox2-mediated metabolism of AA- d8 in the 15 day senescent cells till the end of the experiment (0.25 hour, 0.5 hour, and 1 hour). Senescent MCF7 cells pretreated with Cay-10404 and treated with Cay-10404 and AA-d8 were used as control sample to measure intracellular

metabolism of AA-d8 by Cox2.

- *In vitro* metabolism of AA-d8:
- `o` AA-d8 (250 μM) was incubated with Cox2 enzyme in reaction buffer (100 mM

Tris-Cl (pH=8), 5 mM EDTA, Phenol (500 μM), Hematin (100 nM)).

`o` The reaction was aliquoted in 100 μl aliquots and the aliquots were incubated @37°C till the end of the experiment (0 hour, 0.25 hour, 0.5 hour, 1 hour).
`o` Aliquots were harvested at the end of the experiment and were flash frozen in liquid N2 and were stored @ -80°C.

### Immunofluorescence

- Proliferating and senescent MCF7 cells (Day 10, 15, 18, and 25) were fixed with 4% paraformaldehyde in 1x DPBS @ room temperature for 5 min.
- The cells were washed thrice with 1x DPBS.
- The cells were permeabilized with permeabilization buffer (0.1% Triton X-100 in 1xDPBS) @ room temperature for 15 min.
- The cells were then blocked with blocking buffer (2% Bovine Serum Albumin (BSA) and 2% FBS in the permeabilization buffer) @ room temperature for 90 min.
- The cells were incubated with α-p21 antibody (mouse monoclonal) (Santacruz Biotechnologies) in blocking buffer @ 4°C overnight.
- The cells were washed three times with permeabilization buffer.
- The cells were incubated in Alexa Fluor 568-tagged α-mouse antibody (Cell Signaling Technology) in blocking buffer @ room temperature for 90 min.
- The cells were incubated with DAPI (1 μg/ml) in permeabilization buffer @ room temperature for 10 min.

- The cells were washed thrice with permeabilization buffer.
- The cells were mounted in Prolong Gold antifade medium (Invitrogen).
- The cells were imaged under the FV 3000 inverted microscope (Evident) using appropriate lasers and detectors.

### **X-** gal staining

- Proliferating and senescent MCF7 cells (Day 10, 15, 18, and 25) were fixed with 0.25% glutaraldehyde @ room temperature for 5 min.
- The cells were washed with 1xDPBS.
- The cells were incubated with X-gal staining solution (X-gal (Roche) (1 mg/ml) in 40 mM Citrate-Phosphate Buffer (pH=6), Potassium Ferrocyanide (5 mM), Potassium Ferricyanide (5 mM), Sodium Chloride (150 nM), Magnesium Chloride (2 mM)) @ 37°C overnight.
- The cells were imaged under a Ti2 widefield microscope (Nikon) in brightfield mode.

### Hyperspectral Raman Imaging

- **Label-free:**
- `o` Proliferating and senescent MCF7 cells were fixed with 4% paraformaldehyde @ room temperature for 5 min.
- `o` The cells were washed thrice with 1x DPBS.
- `o` The cells were kept in 1x DPBS during the hyperspectral Raman Imaging.
- `o` Hyperspectral Raman Imaging was done using the alpha 300 Ri system (WITec GmbH, Oxford Instruments) using the following parameters:

♣ Objective: 40x

♣ Laser: 532 nm

♣ Laser power: 60 mW

♣ Scan speed: 2 seconds/pixel

♣ Pixel Size: 1 μm/pixel

♣ Detector Grating: 600 mm^-1^

• AA-d8 labelled:

`o` 15 day Senescent MCF7 cells treated with AA-d8 were fixed with paraformaldehyde @ room temperature for 5 min.

`o` The cells were washed thrice with 1x DPBS.

`o` The cells were kept in 1x DPBS during the hyperspectral Raman Imaging.

`o` Hyperspectral Raman Imaging was done using the alpha 300 Ri system (WITec GmbH, Oxford Instruments) using the following parameters:

♣ Objective: 40x

♣ Laser: 532 nm

♣ Laser power: 60 mW

♣ Scan speed: 2 seconds/pixel

♣ Pixel Size: 1 μm/pixel

♣ Detector Grating: 600 mm^-1^

• Cox2 mediated *in vitro* metabolism of AA-d8.

`o` Flash-frozen samples of Cox2-mediated *in vitro* metabolism of AA-d8 were thawed.

`o` Lipid metabolites were isolated using the Methanol-Formic Acid-Ethyl Acetate method^14,22^.

`o` The samples were resuspended in Ethyl Acetate and were spotted on glass coverslips and dried.

`o` Raman spectra were collected using the alpha 300 Ri system (WITec GmbH) using the following parameters:

♣ Objective: 40x

♣ Laser: 532 nm

♣ Laser power: 60 mW

♣ Acquisition time: 25 seconds/spectrum

♣ Detector Grating: 600 mm^-1^

### Image Analysis

- **True Component Analysis to obtain lipid-rich regions of the Hyperspectral Raman Images:**
- `o` Hyperspectral Raman images were processed in the Project Plus 6.2 (WITec GmbH).
- `o` The images were pre-processed for cosmic ray removal and background subtraction.
- `o` Using the True Component Analysis Pipeline (project plus, WITec GmbH), processed Hyperspectral Raman Images were split into 3 separate components based on spectral de-mixing, and lipid-rich masks were generated (Supplementary Figure S2A).
- `o` The lipid-rich masks were used to generate Hyperspectral Raman Images from the lipid-rich regions of the cells.

### • Spectral Shifts in the deuteration peaks

`o` Raman spectra collected from *in vitro* samples of Cox2-mediated metabolism of AA-d8 were exported as .mat files using Project Plus 6.2 (WITec GmbH).
`o` Spectral analysis was performed using the RamanSPy package^23^ in Python (Appendix I), where:

♣ Raman spectra in the region 2200-2300 cm^-1^ were cropped.

♣ A bimodal curve fitting was done for the deuterium spectra.

♣ Modal values of the fitted curve were obtained to measure spectral shift in the deuterium peaks after in vitro Cox2-mediated metabolism of AA- d8.

• Peak Intensity measurements:

• Hyperspectral Raman Images obtained from proliferating/senescent MCF7 cells or the lipid-rich regions of proliferating/senescent MCF7 cells were exported as

.mat files using Project Plus 6.2 (WITec GmbH).

• The images were processed and analyzed using the RamanSPy package^23^ in Python (Appendix II and III), where:

♣ Raman spectra in the region 1275-1775 cm^-1^ and 2700-3000 cm^-1^ were cropped and concatenated.

♣ Cosmic ray removal was performed using Whitaker-Hayes de-spiking.

♣ Denoising was done using the Savitzky-Golay filter.

• PCA:

♣ Baseline correction was done using the AIRPLS method.

♣ Peak intensities for Raman shifts 1740 cm^-1^ (Ester bond Stretching), 1655 cm^-1^ (C=C Stretching or N-C=O Stretching), 2850 cm^-1^ (CH2 Stretching) 2885 cm^-1^ (CH3 Stretching), and 2920 cm^-1^ (CH3 Stretching) were measured^24,25^.

• Hyperspectral Raman Images obtained from proliferating/senescent MCF7 cells or the lipid-rich regions of proliferating/senescent MCF7 cells were exported as

.mat files using Project Plus 6.2 (WITec GmbH).

• The images were processed and analyzed using the RamanSPy package^23^ in Python (Appendix IV and V), where:

♣ Raman spectra in the region 1275-1775 cm^-1^ and 2700-3000 cm^-1^ were cropped and concatenated.

♣ Cosmic ray removal was performed using Whitaker-Hayes de-spiking.

♣ Denoising was done using the Savitzky-Golay filter.

♣ Baseline correction was done using the AIRPLS method.

♣ Normalization was done using the vector normalization method.

`o` Spectral data from all 5 datasets (Proliferating cells, 10 day senescent cells, 15 day senescent cells, 18 day senescent cells, 25 day senescent cells) were subjected to principal component analysis, where the Raman spectrum from each pixel was considered individually.

`o` Population pairs were plotted on the PC2 vs PC1 plot to visualize population overlaps and differences.

`o` Scatter plot projecting the loadings of the spectral shifts was plotted to visualize the contribution of different spectral shifts to the principal components.

`o` Centroids of the populations were calculated to measure the Euclidean distance between the centroid of given population pair on the PCA plot as a measure of differences in the populations.

• Intracellular Arachidonic Acid Metabolism:

`o` Hyperspectral Raman Images of MCF7 cells treated with AA-d8 were processed in Project Plus 6.2 (WITec GmbH) for Cosmic Ray Removal and Baseline Correction.

`o` Spectral heatmaps of the integrated areas for the peaks corresponding to (C=C)- D Stretching (2220 ± 17.5 cm^-1^ and 2254 ± 17.5 cm^-1^) and CH2 Stretching (2850 ± 15 cm^-1^) were obtained by creating area under the curve filters in Project Plus 6.2 (WITec GmbH).

`o` Ratiometric image analysis was done in ImageJ (NIH) by dividing the spatial heatmaps of (C=C)-D Stretching (2220 ± 17.5 cm^-1^ and 2254 ± 17.5 cm^-1^) by CH2 stretching heatmap (2850 ± 15 cm^-1^) to obtain normalized heatmaps of (C=C)-D stretching peaks to the CH2 stretching peak.

`o` Cell-averaged intensity for the normalized heatmap of (C=C)-D stretching peaks was calculated.

## Results and Discussion

Treatment with DNA-damaging compounds such as Doxorubicin (Doxo) has been shown to induce senescence in cancerous as well as non-cancerous cells^26–28^, including MCF7 human breast adenocarcinoma cells^29,30^. We induced senescence in MCF7 cells by treatment with Doxo (150 nM) for 5 days, followed by culture in DMEM complete medium. The induction of senescence was confirmed by monitoring known markers of senescence, including flattened cell morphology, increased nuclear size, increased expression of tumor suppressor protein p21, and senescence-associated β-galactosidase activity (SA β-gal)^19^. We observed a flattening of the cell morphology with an increase in the cell size (Supplementary Figure S1A), expression of senescence-associated β-galactosidase (SA β-gal) (Supplementary Figure S1B), increased expression of tumor suppressor p21 (Supplementary Figure S1C), and an increase in the nuclear area (Supplementary Figure S1D) in MCF7 cells after treatment with Doxo, confirming the induction of senescence in MCF7 cells upon Doxo mediated DNA damage.

Cells, upon induction of senescence, show an altered metabolic state and acquire a secretory phenotype (Senescence Associated Secretory Phenotype, SASP) and secrete a variety of bioactive molecules, including lipid derivatives^1,11,31–33^. We wanted to capture the temporal dynamics of the altered metabolic states on a single-cell level during senescence. Raman effect-based microscopy allows spatial detection of biomolecular species using characteristic inelastic scattering signatures of biomolecules without extensive sample preparation and labeling^24^. Therefore, we used hyperspectral Raman imaging to analyze intracellular biochemical composition and measure biomolecular distinction between different cell states during senescence. We induced senescence in MCF7 human breast adenocarcinoma cells by treatment with Doxo and fixed the cells with 4% paraformaldehyde on Days 10, 15, 18, and 25 after treatment with Doxo. We used MCF7 cells without treatment with Doxo as a proliferating control. We performed hyperspectral Raman imaging using high laser power (60 mW for 2 seconds/pixel) to achieve an optimal signal-to-noise ratio (∼50 in proliferating cells and >100 in senescent cells) in the Raman spectra. We obtained pixel-by-pixel Raman spectra of proliferating cells and cells on days 10, 15, 18, and 25 after treatment with Doxo. Pre-processing of the hyperspectral Raman images (Denoising, Cosmic Ray Removal, Baseline Correction, Normalization) was done in Python using the RamanSPy package^23^, followed by Principal Component Analysis (PCA) to compare the hyperspectral Raman images of MCF7 cells during senescence (Appendix IV). Visualization of the PCA showed overlap between the Raman spectra of proliferating cells with those of 10 day (Figure 1A), whereas the divergence became apparent by day 15 (Figure 1B), which increased by days 18, and 25 (Figure 1C and D). We also calculated the Euclidean distance between the populations in the PCA space and found that there is a time-dependent increase in the Euclidean distance of the population centroids from the proliferating cells (Supplementary Table 1). We also observed that there is a significant increase in the Euclidean distance between the Raman spectra of proliferating cells and 15 day senescent cells (0.1105 AU) compared to that of proliferating cells and 10 day senescent cells (0.025 AU), suggesting that there is a major change in the Raman signatures of MCF7 between 10 and 15 days after treatment with Doxo.

**Figure 1:**
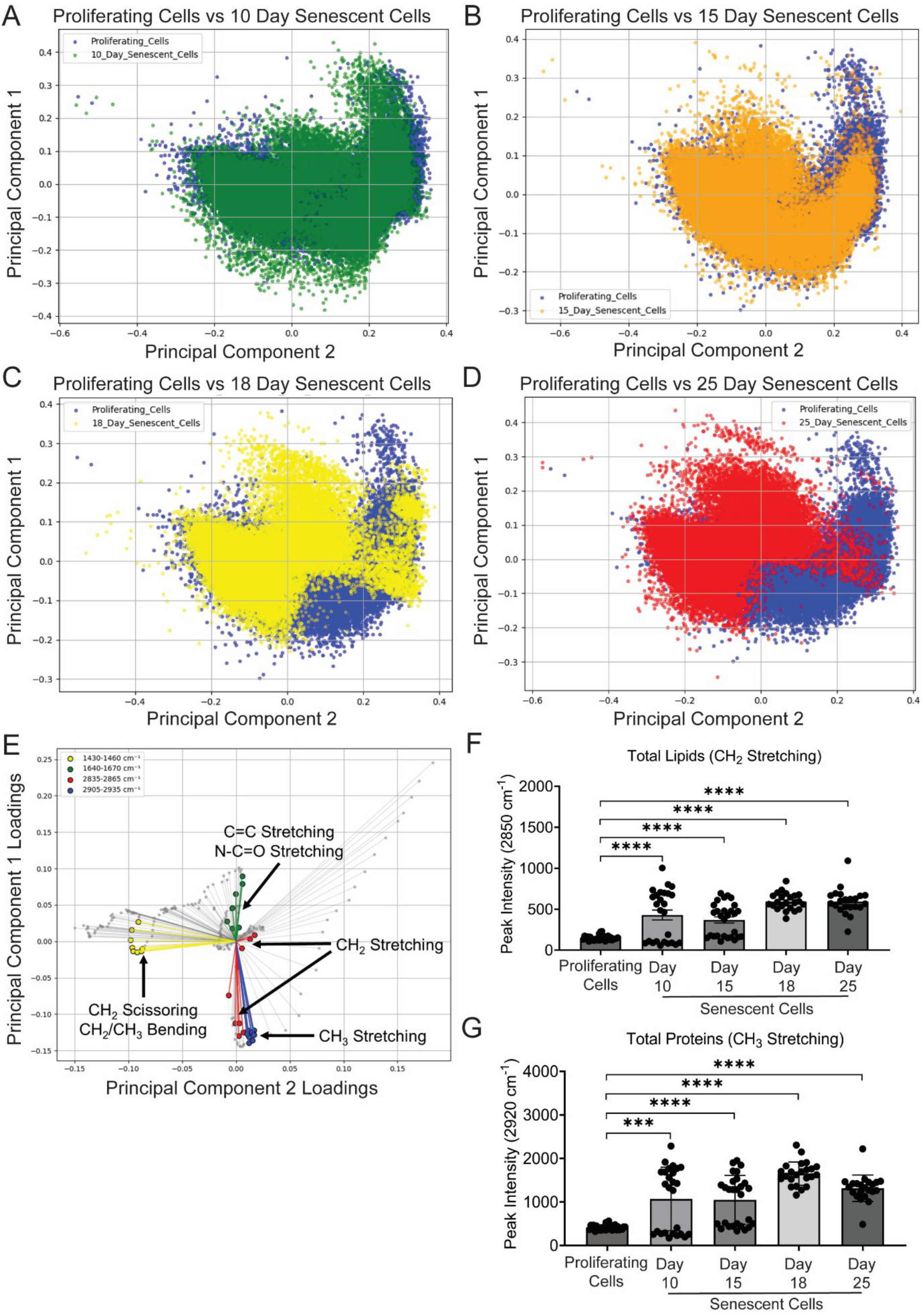
Visualization of intracellular biomolecular composition of senescent cells using Hyperspectral Raman Imaging. A. Principal Component Analysis of Hyperspectral Raman Images of MCF7 human breast adenocarcinoma cells during proliferation and 10 days after induction of DNA damage- mediated senescence (n ≥ 20 cells). B. Principal Component Analysis of Hyperspectral Raman Images of MCF7 human breast adenocarcinoma cells during proliferation and 15 days after induction of DNA damage- mediated senescence (n ≥ 20 cells). C. Principal Component Analysis of Hyperspectral Raman Images of MCF7 human breast adenocarcinoma cells during proliferation and 18 days after induction of DNA damage- mediated senescence (n ≥ 20 cells). D. Principal Component Analysis of Hyperspectral Raman Images of MCF7 human breast adenocarcinoma cells during proliferation and 25 days after induction of DNA damage- mediated senescence (n ≥ 20 cells). E. Scatter plot of the projected loadings of the Raman shifts’ contribution to the principal components PC1 and PC2 in the PCA of hyperspectral Raman images of MCF7 human breast adenocarcinoma cells during DNA damage-mediated senescence after treatment with Doxo (150 nM). Red highlighted points show projected loadings of Raman shifts corresponding to CH2 stretching (2850 cm^-1^), green points show projected loadings of Raman shifts corresponding to C=C or N-C=O Stretching (1655 cm^-1^), blue points show projected loadings of Raman shifts corresponding to CH3 stretching (2920 cm^-1^), and the yellow points show projected loadings of Raman shifts corresponding to CH2 scissoring or CH2/CH3 bending (1445 cm^-1^). F. Cell-averaged peak intensity of the CH2 stretching peak (2850 cm^-1^) obtained from hyperspectral Raman Imaging of MCF7 human breast adenocarcinoma cells during DNA damage-mediated senescence (n ≥ 20 cells). G. Cell-averaged peak intensity of the CH3 stretching peak (2920 cm^-1^) obtained from hyperspectral Raman Imaging of MCF7 human breast adenocarcinoma cells during DNA damage-mediated senescence (n ≥ 20 cells). (The standard deviation between replicates was plotted as error bars. Statistical significance was tested by the two-tailed Student’s t-test assuming heteroscedastic distributions. ***: p<0.001, ****: p<0.0001)

Spectral loadings of the PCA demonstrated that lipid-related vibrational modes contributed to the separation of populations on the PCs (Figure 1E): CH2 scissoring (1445 cm^-1^) accounts for ∼0.0716, C=C stretching (1655 cm^-1^) contributed ∼0.0223, and CH2 stretching (2850 cm^-1^) contributed ∼ 0.065 to the PCs. Also, measuring the intensities of spectral shift peaks corresponding to biomolecular species showed a time-dependent increase in the spectral peak corresponding to CH2 stretching (2850 cm^-1^) (Figure 1F). Spectral peak intensity of C=C/N-C=O Stretching (1655 cm^-1^) showed a time-dependent increase till day 18, followed by a decrease in intensity by day 25 (Supplementary Figure S1E). Spectral peak intensity of CH3 Stretching (2920 cm^-1^ and 2885 cm^-1^) also showed a time-dependent increase till day 18, followed by a decrease in intensity by day 25 (Figure 1G and Supplementary Figure S1F). Finally, the spectral peak intensity of ester bond stretching (1740 cm^-1^) corresponding to triglycerides showed a time-dependent increase in senescent MCF7 cells (Supplementary Figure S1G). This suggests that there is a distinct time-dependent alteration of metabolism of different biomolecular species during senescence.

Storage and metabolism of fatty acids in cells are compartmentalized in the cells^34,35^. We used hyperspectral Raman imaging of fixed cells, followed by true component analysis (WITec GmbH) (Supplementary Figure S2A), to obtain Raman spectra from the lipid-rich regions in MCF7 cells during senescence (Figure 2A). We obtained hyperspectral Raman images from the lipid-rich regions of proliferating MCF7 cells and of those on 10, 15, 18, and 25 days after treatment with Doxo, pre-processed the data in Python using RamanSPy^23^, and performed PCA on the processed data (Appendix V). We observed an overlap between the Raman spectra of the proliferating cells vs 10 day senescent cells (Figure 2B), and the divergence became apparent from day 15 (Figure 2C), which increased on days 18 and 25 (Figure 2D and E), similar to the PCA of whole cells (Figure 1A, B, C, and D). Calculation of the Euclidean distance between the population showed that there is a time-dependent increase in the Euclidean distance of the population centroids from the proliferating cells (Supplementary Table 2). We also observed a significant increase in the Euclidean distance between the Raman spectra of proliferating cells and 15 day senescent cells (0.0905) compared to that of proliferating cells and 15 day senescent cells (0.0396), suggesting that there is a major change in the Raman signatures of lipid-rich regions of MCF7 cells between 10 and 15 days after treatment with Doxo. Analysis of the spectral loadings of the PCA demonstrated that lipid-related vibrational modes contributed to the separation of populations on the PCs (Figure 2F): CH2 scissoring (1445 cm^-1^) accounts for ∼0.2624, C=C stretching (1655 cm^-1^) contributed ∼0.2413, and CH2 stretching (2850 cm^-1^) contributed ∼ 1.142. Measuring the intensities of the spectral shift peak corresponding to CH2 stretching (2850 cm^-1^), CH3 Stretching (2920 cm^-1^ and 2885 cm^-1^), C=C/N-C=O Stretching (1655 cm^-1^), and ester bond stretching (1740 cm^-1^) showed an increase in the intensity till day 18, with day 18 and 25 showing similar intensities (Figure 2G, Supplementary Figures S2B, C, D, and E). These findings indicate that fatty acid– associated spectral shifts underlie the differentiation of proliferating and senescent cell states by hyperspectral Raman imaging. Together, the results establish that Raman microscopy, coupled with True Component Analysis and PCA, can resolve senescence-associated lipid remodeling with high specificity. This capability highlights the potential to identify senescent cell niches in tissues in a label-free manner, enabling strategies for selective targeting of senescent cells in aging and disease.

**Figure 2:**
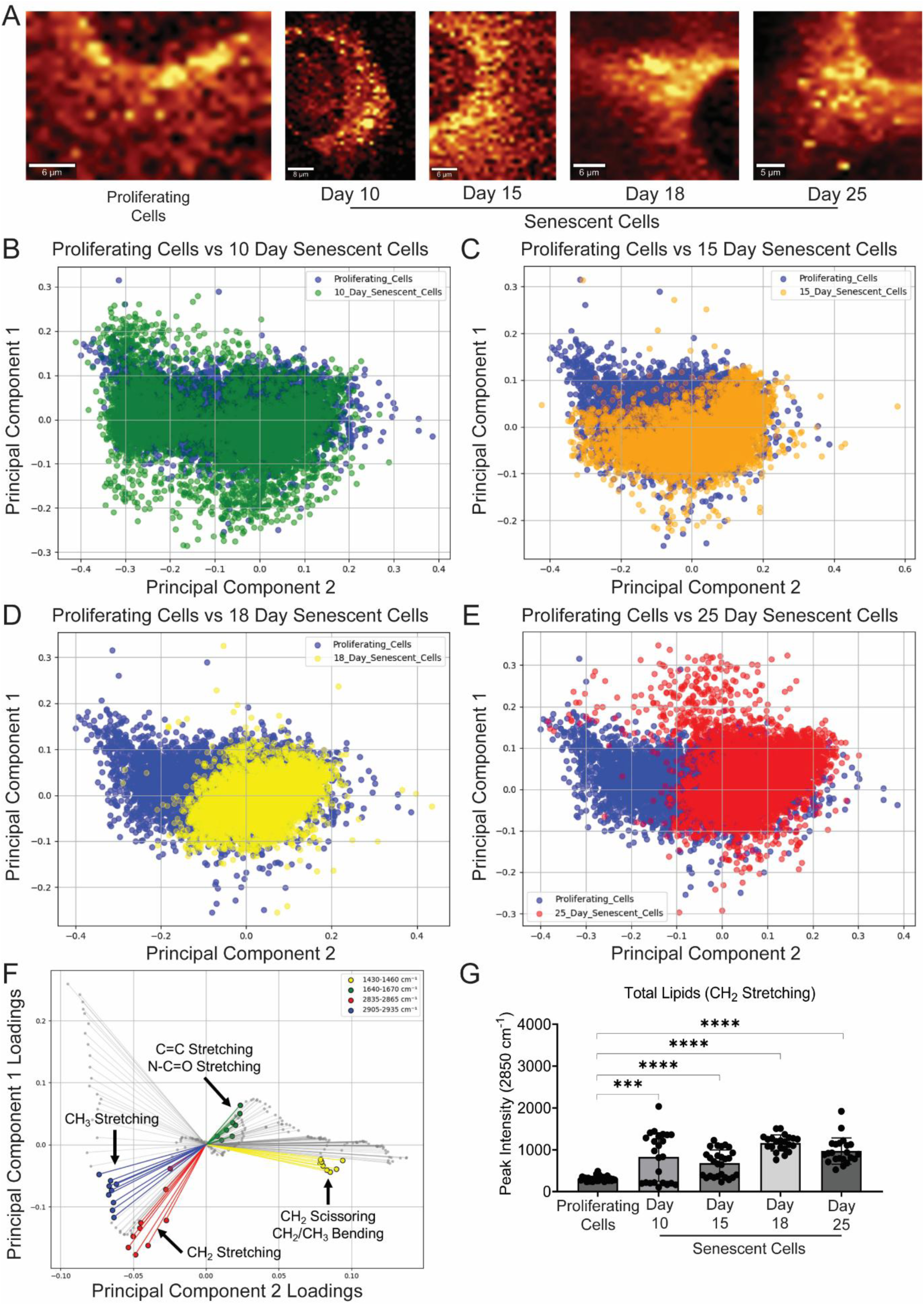
Visualization of intracellular biomolecular composition of lipid-rich regions in senescent cells using Hyperspectral Raman Imaging. A. Representative heatmaps of lipid-rich regions in MCF7 human breast adenocarcinoma cells during DNA damage-mediated senescence after treatment with Doxo (150 nM). B. Principal Component Analysis of Raman spectra isolated from the lipid-rich regions of MCF7 human breast adenocarcinoma cells during proliferation and 10 days after induction of DNA damage-mediated senescence (n ≥ 20 cells). C. Principal Component Analysis of Raman spectra isolated from the lipid-rich regions of MCF7 human breast adenocarcinoma cells during proliferation and 15 days after induction of DNA damage-mediated senescence (n ≥ 20 cells). D. Principal Component Analysis of Raman spectra isolated from the lipid-rich regions of MCF7 human breast adenocarcinoma cells during proliferation and 18 days after induction of DNA damage-mediated senescence (n ≥ 20 cells). E. Principal Component Analysis of Raman spectra isolated from the lipid-rich regions of MCF7 human breast adenocarcinoma cells during proliferation and 25 days after induction of DNA damage-mediated senescence (n ≥ 20 cells). F. Scatter plot of the projected loadings of the Raman shifts’ contribution to the principal components PC1 and PC2 in the PCA of pixel-by-pixel Raman spectra obtained from the lipid-rich regions of MCF7 human breast adenocarcinoma cells during DNA damage- mediated senescence after treatment with Doxo (150 nM). Red highlighted points show projected loadings of Raman shifts corresponding to CH2 stretching (2850 cm^-1^), green points show projected loadings of Raman shifts corresponding to C=C or N-C=O Stretching (1655 cm^-1^), blue points show projected loadings of Raman shifts corresponding to CH3 stretching (2920 cm^-1^), and the yellow points show projected loadings of Raman shifts corresponding to CH2 scissoring or CH2/CH3 bending (1445 cm^-1^). G. Averaged peak intensity of the CH2 stretching peak (2850 cm^-1^) in Raman spectra obtained from the lipid-rich regions of MCF7 human breast adenocarcinoma cells during DNA damage-mediated senescence (n ≥ 20 cells). (The standard deviation between replicates was plotted as error bars. Statistical significance was tested by the two-tailed Student’s t-test assuming heteroscedastic distributions. ***: p<0.001, ****: p<0.0001)

Senescent cells show increased accumulation of Polyunsaturated Fatty Acids (PUFA), including Arachidonic Acid^11^. Senescent cells also show increased metabolism of Arachidonic Acid, leading to synthesis and release of several classes of oxylipins, including prostaglandins^11,14^. Therefore, we measured the metabolism of arachidonic acid in senescent cells using Raman microscopy. Arachidonic Acid (20:4) is a polyunsaturated fatty acid that shows characteristic spectral shifts at wave numbers 1655 cm^-1^ and 3005 cm^-1^ that correspond to C=C stretching and (C=C)-H stretching, respectively (Figure S3A). Arachidonic Acid with deuterium atoms across all the double bonds shows the presence of a characteristic spectral shift corresponding to (C=C)-D stretching at ∼2240 cm^-1^ (Figure S3A, Figure 3A) in the biologically silent region^36^. Measuring the changes in the (C=C)-D stretching peaks is advantageous due to the high signal-to-noise ratio in the otherwise silent region, allowing measurement of chemical changes only in the labelled molecule. Therefore, we used deuterated Arachidonic Acid (AA-d8) to measure the intracellular metabolism of Arachidonic Acid in senescent cells. We observed that the Raman spectrum of AA-d8 shows two peaks corresponding to (C=C)-D stretching, one at 2220 cm^-1^ (minor peak) and another at 2254 cm^-1^ (major peak), suggesting two vibrational modes of (C=C)-D stretching (Figure 3A). We hypothesized that the metabolism of arachidonic acid can be measured by changes in the vibrational modes of the (C=C)-D bond, which can be monitored by changes in the Raman signatures. We tested the hypothesis by *in vitro* metabolism of AA-d8 by the COX2 enzyme (Figure S3B) and measuring the changes in the spectral shifts. We did not observe any change in the wavenumber of the spectral shifts in the (C=C)-D peaks (Figure S3C), suggesting no change in the vibrational modes of (C=C)-D stretching. We then measured any changes in the relative intensities of the (C=C)-D peaks (2220 cm^-1^ and 2254 cm^-1^) to the CH2 stretching peaks (2850 cm^-1^) during COX2-mediated metabolism of arachidonic acid. We observed a time-dependent decrease in the intensities of the (C=C)-D peaks relative to the CH2 Stretching peaks during COX2- mediated metabolism of AA-d8, suggesting that the relative intensities of the (C=C)-D peaks can be used to measure metabolism of Arachidonic Acid. So, to measure the metabolism of arachidonic acid in senescent cells, we treated ∼15-day senescent MCF7 cells with AA-d8 (100 μM) overnight and observed complete cell death (data not shown). We hypothesized that the cell death observed after treating senescent cells with AA-d8 overnight is because of the cytotoxic effects of arachidonic acid metabolites, a phenotype we observed in a previous study^14^. Therefore, we pretreated senescent MCF7 cells with Cay-10404, an inhibitor of COX2, for 2 hours before treating the cells with AA-d8. We treated the pretreated senescent MCF7 cells with AA-d8 (100 μM) in the presence of Cay-10404 overnight. We then removed the Cay-10404 from the treatment medium and incubated the cells with AA-d8 for 0.25, 0.5, and 1 hour. We then fixed the cells and performed hyperspectral Raman imaging. Spectral heatmaps of the senescent MCF7 in the regions 2220±17.5 cm^-1^ and 2254±17.5 cm^-1^ showed intracellular accumulation of AA-d8 in the senescent cells, predominantly in the perinuclear region (Figure 3C). We also observed a heterogeneous intensity distribution of AA-d8 in the perinuclear region, suggesting a heterogeneity in the intracellular accumulation of Arachidonic Acid. We analyzed the hyperspectral Raman images of senescent cells treated with AA-d8 by ratiometric analysis of spectral heatmaps of spectral shifts corresponding to (C=C)-D stretching to CH2 stretching (Figure 3D) and observed a time-dependent decrease in the major peak (2254 cm^-1^) intensity normalized to total lipids (2850 cm^-1^) after removal of COX2 inhibitor (Figure 3E) similar to that in the *in vitro* metabolism of AA-d8 (Figure 3B), suggesting metabolism of arachidonic acid in senescent MCF7 cells. We also observed a decrease in the peak intensity of the minor peak (2220 cm^-1^) normalized to total lipids (2850 cm^-1^) at 1 hour after removal of the COX2 inhibitor. These observations suggest the detection of the metabolism of arachidonic acid in senescent MCF7 cells after removal of the COX2 inhibitor by Raman microscopy. This approach demonstrates that Raman spectroscopy of deuterated arachidonic acid can monitor COX2-driven metabolism both in vitro and in cells. Beyond validating COX2 inhibition, this strategy opens the possibility of screening inhibitors of other enzymes in arachidonic acid pathways. By coupling enzymatic assays with hyperspectral Raman imaging, one can spatially and temporally map lipid metabolism in senescent cells at high resolution. This capacity to resolve metabolic heterogeneity without labels establishes Raman microscopy as a powerful chemical biology tool: enabling precise dissection of senescence-associated lipid pathways and accelerating discovery of selective interventions against senescent cell function.

**Figure 3.**
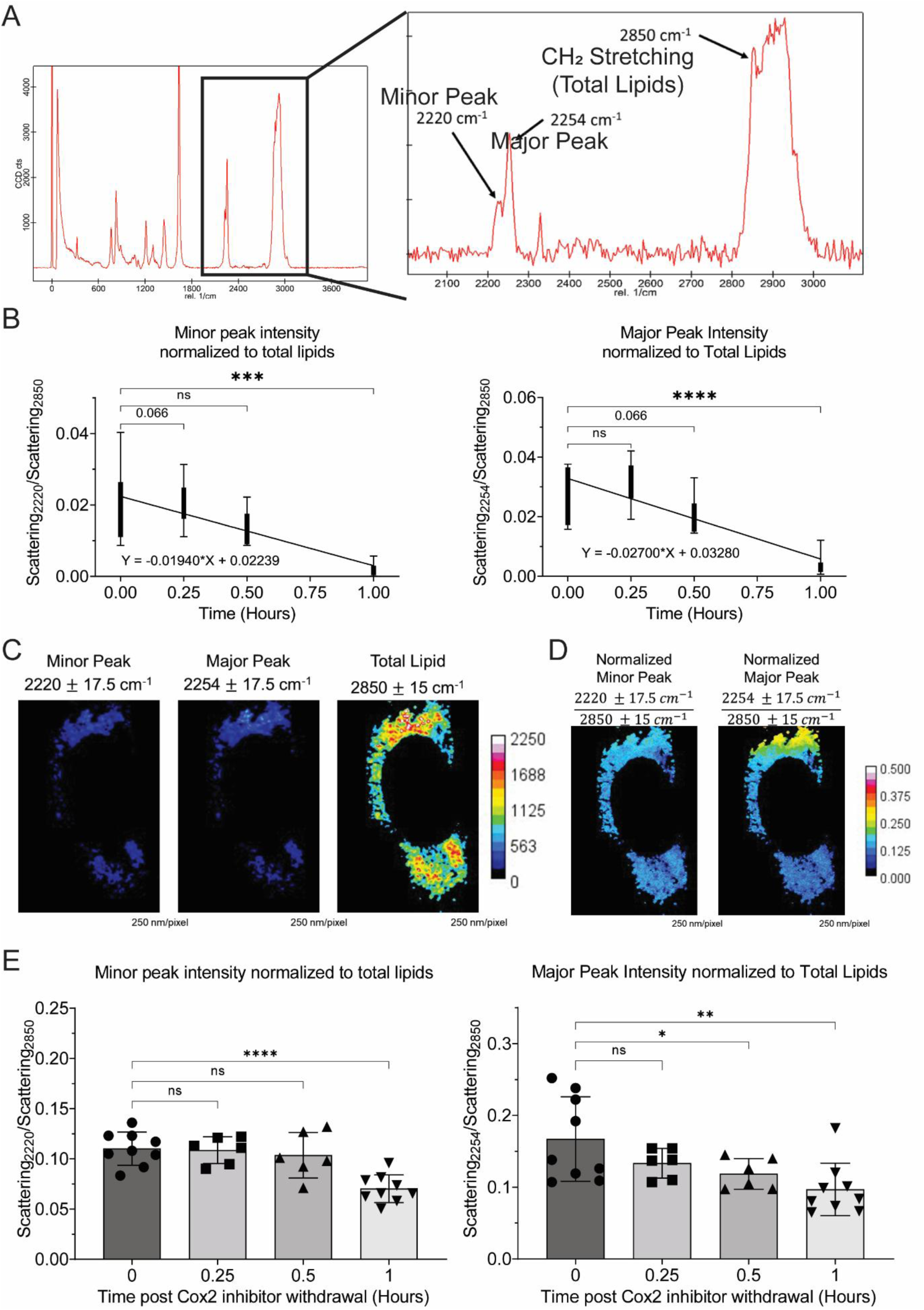
Visualization of Arachidonic Acid Metabolism in senescent MCF7 cells using hyperspectral Raman imaging of deuterated arachidonic acid. A. Raman spectrum of 5,6,8,9,11,12,14,15 D8 – Arachidonic Acid (AA-d8) showing two modes of signature spectral shifts of (C=C)-D stretching in the biologically silent region at 2220 cm^-1^ (minor peak) and 2254 cm^-1^ (major peak). B. Intensity of the two (C=C)-D stretching peaks (2220 and 2254 cm^-1^) normalized to the intensity of the total lipids peak (CH2 Stretching, 2850 cm^-1^) measured by Raman spectroscopy at different timepoints during PTGS2-mediated metabolism of Arachidonic Acid in *vitro* (n≥15). C. Visualization of the intracellular distribution of AA-d8 in senescent MCF7 cells by hyperspectral Raman imaging. D. Ratiometric heatmaps for visualization of the intracellular distribution of AA-d8 in senescent MCF7 cells by hyperspectral Raman imaging. E. Intensity of the two (C=C)-D stretching peaks (2220 and 2254 cm^-1^) normalized to the intensity of the total lipids peak (CH2 Stretching, 2850 cm^-1^) measured by hyperspectral Raman imaging in senescent MCF7 cells at different timepoints after removal of PTGS2 (Cox2) inhibitor (CAY 10404). (The standard deviation between replicates was plotted as error bars. Statistical significance was tested by the two-tailed Student’s t-test assuming heteroscedastic distributions. ***: p<0.001, ****: p<0.0001)

## Supporting Information for publication

Appendix I: Python script to measure the shift in the Raman peaks corresponding to AA-d8 during Cox2-mediated in vitro metabolism.

Appendix II: Python script to measure the intensity of Raman peaks corresponding to different biomolecular species in MCF7 cells during senescence.

Appendix III: Python script to measure the intensity of Raman peaks corresponding to different biomolecular species in the lipid-rich regions of MCF7 cells during senescence.

Appendix IV: Python script for principal component analysis of hyperspectral Raman images of MCF7 cells during senescence.

Appendix V: Python script for principal component analysis of hyperspectral Raman images of lipid-rich regions of MCF7 cells during senescence.

## AUTHOR INFORMATION

Corresponding Author

Dr. Arvind Ramanathan

Metabolic Regulation of Cell Fate (RCF), Institute for Stem Cell Science and Regenerative Medicine (BRIC-InStem), GKVK – Post, Bellary Road, Bengaluru – 560065.

Contact: >

## Author Contributions

SSP and AR conceptualized the project, designed experimental protocols and analysis pipelines, and wrote the manuscript. SSP did the *in vitro* biochemical experiments, Raman Imaging, and Image processing (both ImageJ and RamanSPy). AV performed the cell culture experiments, helped with Image processing (RamanSPy), and manuscript edits. HM helped with reagent procurement and Raman imaging and analysis. AR funded the project and is the corresponding author. The manuscript was written through the contributions of all authors. All authors have given approval to the final version of the manuscript.

## Funding Sources

### ACKNOWLEDGMENT

The authors thank the Central Imaging and Flow Cytometry Facility (NCBS-InStem) for the imaging solutions. The Authors thank Prof. Satyajit Mayor for his valuable inputs in the study.

### ABBREVIATIONS

DNA: Deoxyribonucleic Acid, SASP: Senescence Associated Secretory Phenotype, MCF7: Human Breast Adenocarcinoma Cell Line Michigan Cell Foundation 7, PCA: Principal Component Analysis, AA-d8: 5,6,8,9,11,12,14,15 D8 – Arachidonic Acid, COX2: Cyclooxygenase 2 enzyme, Doxo: Doxorubicin, PUFA: Polyunsaturated Fatty Acids, DMEM: Dulbecco’s Modified Eagle Medium, FBS: Fetal Bovine Serum, DMSO: Dimethyl Sulphoxide, DPBS: Dulbecco’s Phosphate Buffer Saline, EDTA: Ethylenediaminetetraacetic Acid

---

**Supplementary Figure S1:**
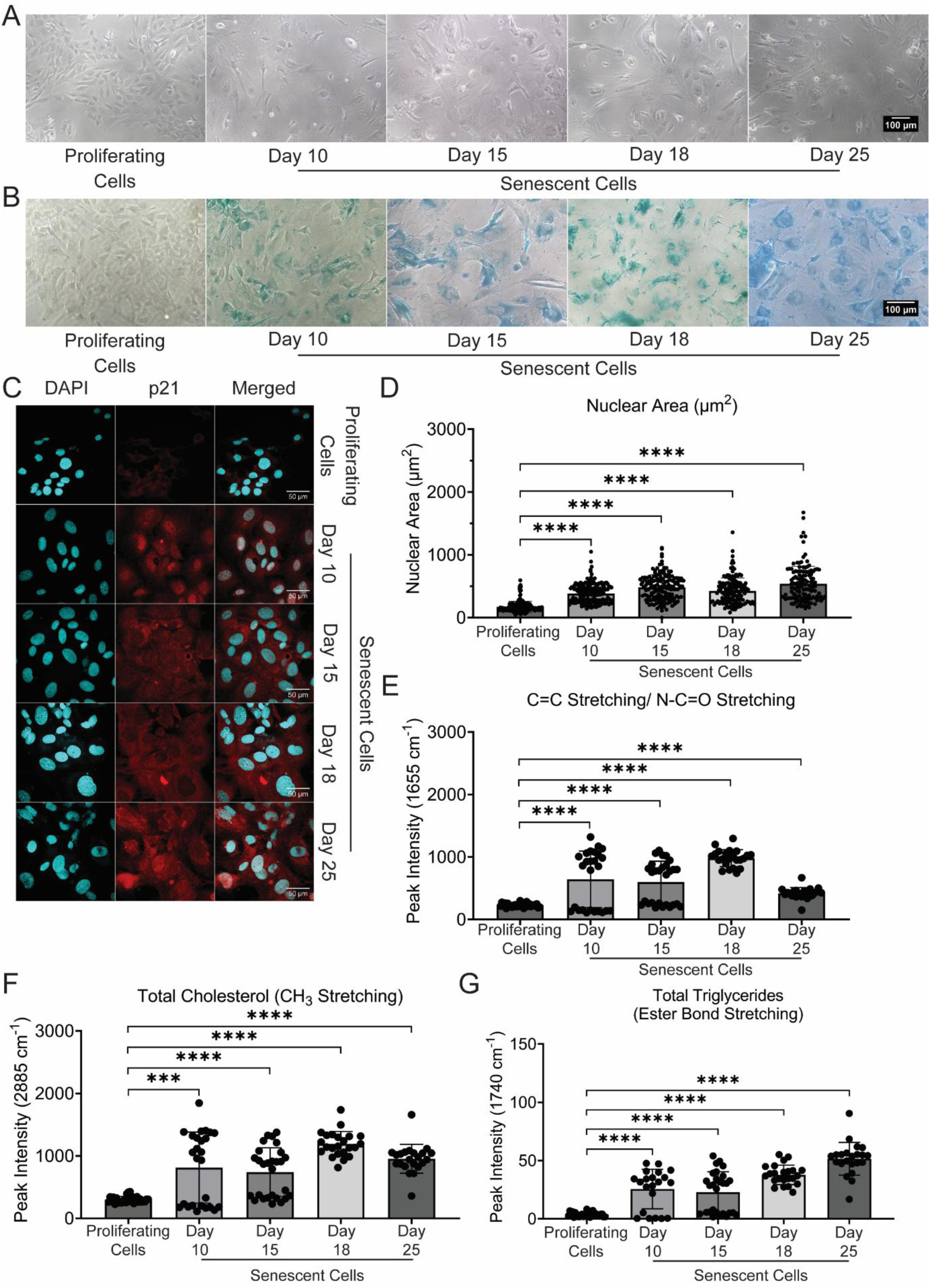
Visualization of intracellular biomolecular composition of senescent cells using Hyperspectral Raman Imaging. A. Morphology of MCF7 human breast adenocarcinoma cells during DNA damage-mediated senescence after treatment with Doxo (150 nM). B. Expression of senescence-associated β-galactosidase (SA β-gal) in MCF7 human breast adenocarcinoma cells during DNA damage-mediated senescence after treatment with Doxo (150 nM). C. Expression and localization of tumor suppressor p21 in MCF7 human breast adenocarcinoma cells after DNA damage-mediated senescence after treatment with Doxo (150 nM). D. Nuclear size in MCF7 human breast adenocarcinoma cells after DNA damage-mediated senescence after treatment with Doxo (150 nM). E. Cell-averaged peak intensity of the C=C/N-C=O stretching peak (1655 cm^-1^) obtained from hyperspectral Raman Imaging of MCF7 human breast adenocarcinoma cells during DNA damage-mediated senescence (n ≥ 20 cells). F. Cell-averaged peak intensity of the CH3 stretching peak (2885 cm^-1^) obtained from hyperspectral Raman Imaging of MCF7 human breast adenocarcinoma cells during DNA damage-mediated senescence (n ≥ 20 cells). G. Cell-averaged peak intensity of the ester bond stretching (1740 cm^-1^) obtained from hyperspectral Raman Imaging of MCF7 human breast adenocarcinoma cells during DNA damage-mediated senescence (n ≥ 20 cells). (The standard deviation between replicates was plotted as error bars. Statistical significance was tested by the two-tailed Student’s t-test assuming heteroscedastic distributions. ***: p<0.001, ****: p<0.0001)

**Supplementary Figure S2:**
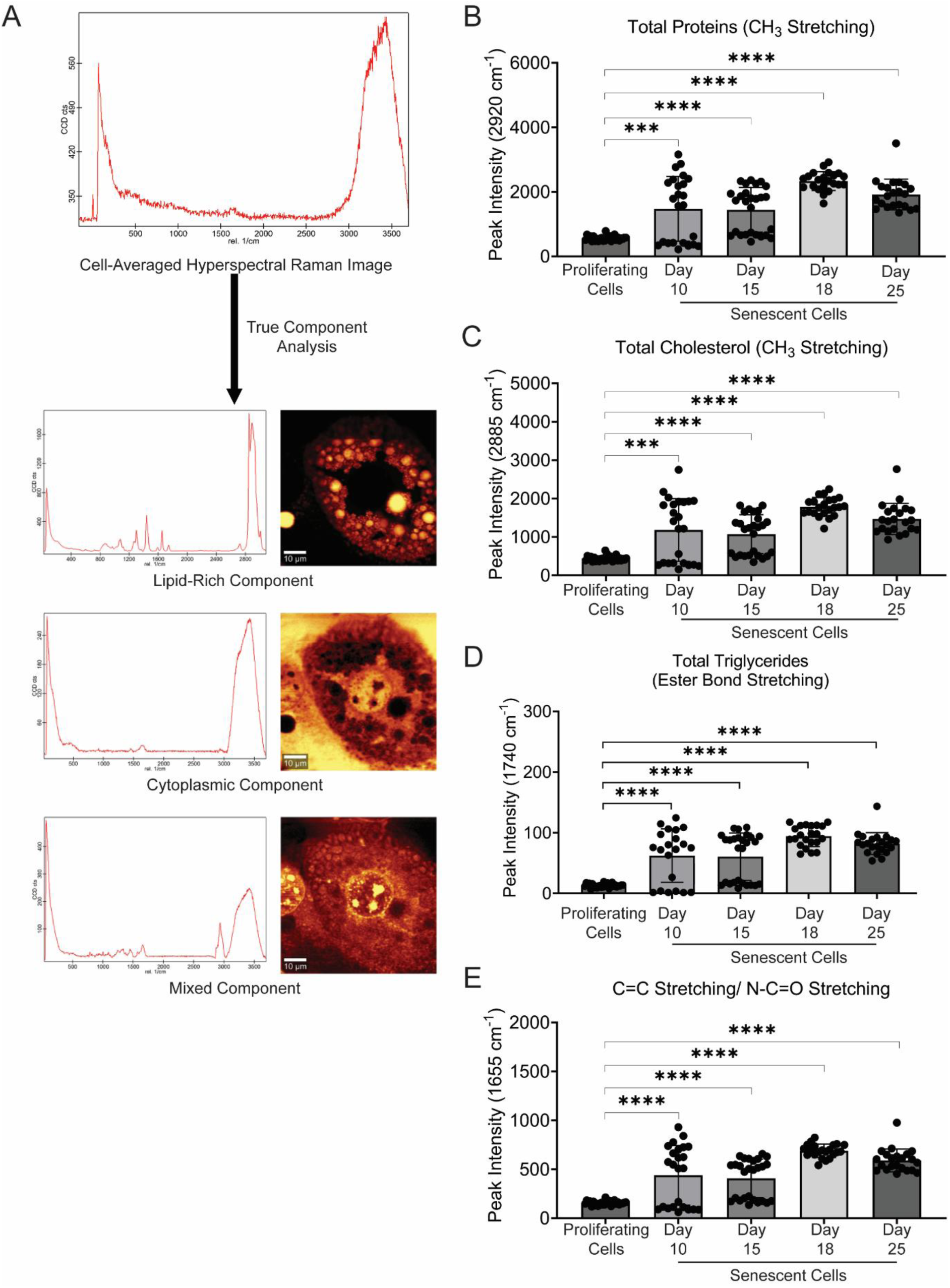
Visualization of intracellular biomolecular composition of lipid-rich regions in senescent cells using Hyperspectral Raman Imaging. A. Schematic representation of the True Component Analysis (WITec Project 6.2), and the resultant spatial heatmaps and representative Raman spectra, from lipid-rich regions, cytoplasmic region, and region with mixed Raman signatures. B. Averaged peak intensity of the CH3 stretching peak (2920 cm^-1^) in Raman spectra obtained from the lipid-rich regions of MCF7 human breast adenocarcinoma cells during DNA damage-mediated senescence (n ≥ 20 cells). C. Averaged peak intensity of the CH3 stretching peak (2885 cm^-1^) in Raman spectra obtained from the lipid-rich regions of MCF7 human breast adenocarcinoma cells during DNA damage-mediated senescence (n ≥ 20 cells). D. Cell-averaged peak intensity of the Ester bond stretching (1740 cm^-1^) obtained from the lipid-rich regions of MCF7 human breast adenocarcinoma cells during DNA damage-mediated senescence (n ≥ 20 cells). E. Cell-averaged peak intensity of the C=C/N-C=O stretching peak (1655 cm^-1^) obtained from the lipid-rich regions of MCF7 human breast adenocarcinoma cells during DNA damage-mediated senescence (n ≥ 20 cells). (The standard deviation between replicates was plotted as error bars. Statistical significance was tested by the two-tailed Student’s t-test assuming heteroscedastic distributions. ***: p<0.001, ****: p<0.0001)

**Supplementary Figure S3:**
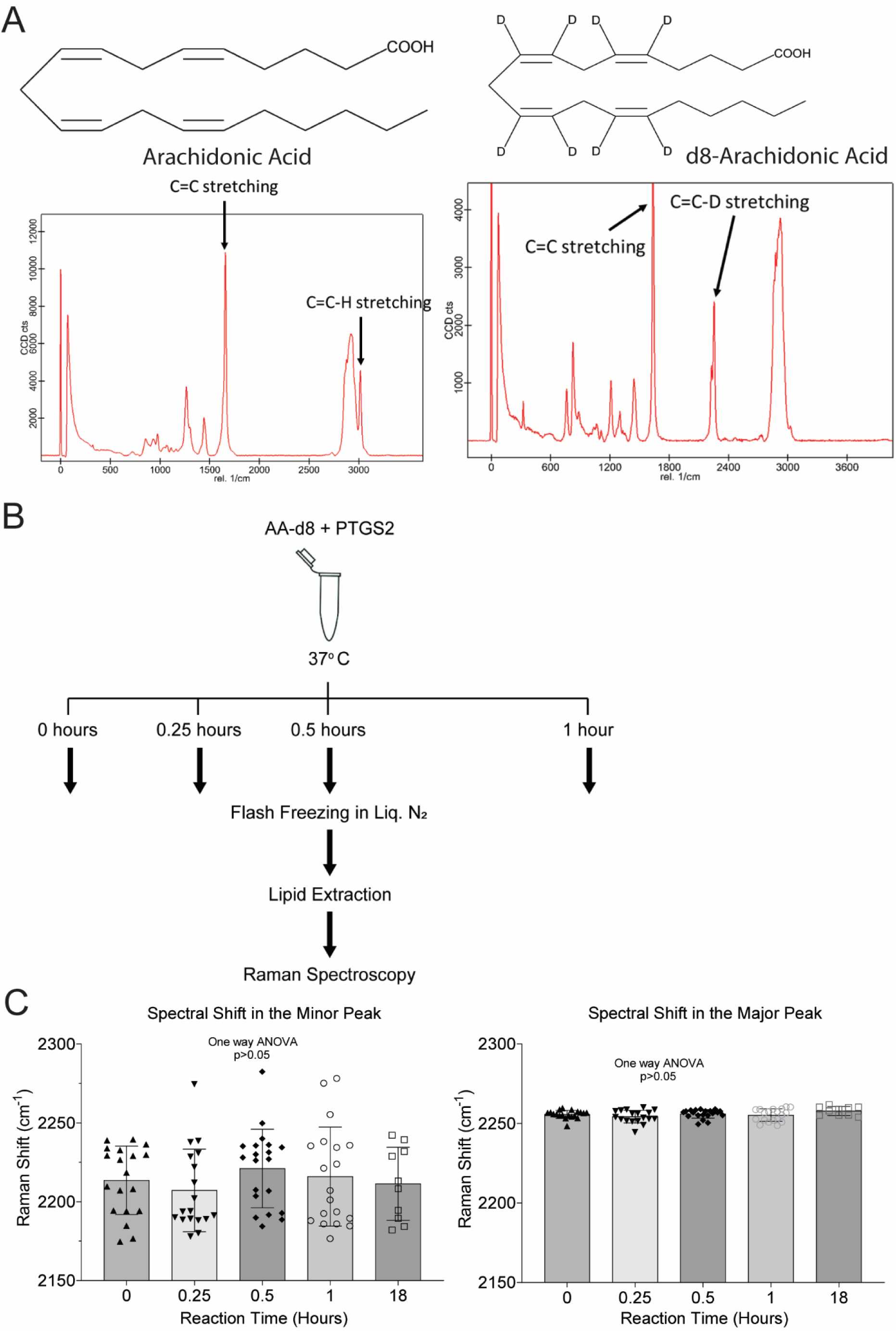
Visualization of Arachidonic Acid Metabolism using hyperspectral Raman imaging of deuterated arachidonic acid. A. Raman spectra of Arachidonic Acid and AA-d8 showing the presence of spectral shifts corresponding to (C=C)-H stretching (3005 cm^-1^) and (C=C)-D Stretching (2220 and 2254 cm^-1^), respectively. B. Schematic Representation of the PTGS2-mediated metabolism of AA-d8 *in vitro*. C. Spectral shift wavenumbers corresponding to (C=C)-D Stretching in AA-d8 at different timepoints during PTGS2-mediated metabolism *in vitro*.

**Supplementary Table 1:** Euclidean distance between the centroids of the population pairs on the PCA plot of hyperspectral Raman images of MCF7 cells during senescence.

| **Population Pair** | **Euclidean Distance between Population Centroids on the PCA plot (AU)** |
| --- | --- |
| Proliferating Cells vs 10 Day Senescent Cells | 0.025 |
| Proliferating Cells vs 15 Day Senescent Cells | 0.1105 |
| Proliferating Cells vs 18 Day Senescent Cells | 0.1234 |
| Proliferating Cells vs 25 Day Senescent Cells | 0.1860 |

**Supplementary Table 2:** Euclidean distance between the centroids of the population pairs on the PCA plot of hyperspectral Raman images of lipid-rich regions in MCF7 cells during senescence.

| **Population Pair** | **Euclidean Distance between Population Centroids on the PCA plot (AU)** |
| --- | --- |
| Proliferating Cells vs 10 Day Senescent Cells | 0.0396 |
| Proliferating Cells vs 15 Day Senescent Cells | 0.0905 |
| Proliferating Cells vs 18 Day Senescent Cells | 0.1015 |
| Proliferating Cells vs 25 Day Senescent Cells | 0.1200 |

## REFERENCES

(1) Coppé, J. P.; Desprez, P. Y.; Krtolica, A.; Campisi, J. The Senescence-Associated Secretory Phenotype: The Dark Side of Tumor Suppression. Annu. Rev. Pathol. Mech. Dis. 2010, 5, 99–118. 10.1146/annurev-pathol-121808-102144.

(2) Campisi, J. Aging, Cellular Senescence, and Cancer. Annu. Rev. Physiol. 2013, 75, 685– 705. 10.1146/ANNUREV-PHYSIOL-030212-183653.

(3) Kuilman, T.; Peeper, D. S. Senescence-Messaging Secretome: SMS-Ing Cellular Stress. Nat. Rev. Cancer 2009 92 2009, 9 (2), 81–94. 10.1038/nrc2560.

(4) Di, X.; Bright, A. T.; Bellott, R.; Gaskins, E.; Robert, J.; Holt, S.; Gewirtz, D.; Elmore, L. A Chemotherapy-Associated Senescence Bystander Effect in Breast Cancer Cells. Cancer Biol. Ther. 2008, 7 (6), 864–872. 10.4161/CBT.7.6.5861.

(5) Bojko, A.; Czarnecka-Herok, J.; Charzynska, A.; Dabrowski, M.; Sikora, E. Diversity of the Senescence Phenotype of Cancer Cells Treated with Chemotherapeutic Agents. Cells 2019, 8 (12), 1501. 10.3390/CELLS8121501.

(6) Sikora, E.; Mosieniak, G.; Alicja Sliwinska, M. Morphological and Functional Characteristic of Senescent Cancer Cells. Curr. Drug Targets 2016, 17 (4), 377–387. 10.2174/1389450116666151019094724.

(7) Davalos, A. R.; Coppe, J. P.; Campisi, J.; Desprez, P. Y. Senescent Cells as a Source of Inflammatory Factors for Tumor Progression. Cancer Metastasis Rev. 2010, 29 (2), 273–283. 10.1007/s10555-010-9220-9.

(8) Wiley, C. D.; Campisi, J. From Ancient Pathways to Aging Cells - Connecting Metabolism and Cellular Senescence; Cell Press, 2016; Vol. 23, pp 1013–1021. 10.1016/j.cmet.2016.05.010.

(9) Lizardo, D. Y.; Lin, Y. L.; Gokcumen, O.; Atilla-Gokcumen, G. E. Regulation of Lipids Is Central to Replicative Senescence. Mol. Biosyst. 2017, 13 (3), 498–509. 10.1039/C6MB00842A.

(10) Chong, M.; Yin, T.; Chen, R.; Xiang, H.; Yuan, L.; Ding, Y.; Pan, C. C.; Tang, Z.; Alexander, P. B.; Li, Q.; Wang, X. CD36 Initiates the Secretory Phenotype during the Establishment of Cellular Senescence. EMBO Rep. 2018, 19 (6). 10.15252/EMBR.201745274.

(11) Wiley, C. D.; Sharma, R.; Davis, S. S.; Lopez-Dominguez, J. A.; Mitchell, K. P.; Wiley, S.; Alimirah, F.; Kim, D. E.; Payne, T.; Rosko, A.; Aimontche, E.; Deshpande, S. M.; Neri, F.; Kuehnemann, C.; Demaria, M.; Ramanathan, A.; Campisi, J. Oxylipin Biosynthesis Reinforces Cellular Senescence and Allows Detection of Senolysis. Cell Metab. 2021, 33 (6), 1124–1136.e5. 10.1016/j.cmet.2021.03.008.

(12) Martien, S.; Pluquet, O.; Vercamer, C.; Malaquin, N.; Martin, N.; Gosselin, K.; Pourtier, A.; Abbadie, C. Cellular Senescence Involves an Intracrine Prostaglandin E2 Pathway in Human Fibroblasts. Biochim. Biophys. Acta - Mol. Cell Biol. Lipids 2013, 1831 (7), 1217–1227. 10.1016/j.bbalip.2013.04.005.

(13) Wiley, C. D.; Brumwell, A. N.; Davis, S. S.; Jackson, J. R.; Valdovinos, A.; Calhoun, C.; Alimirah, F.; Castellanos, C. A.; Ruan, R.; Wei, Y.; Chapman, H. A.; Ramanathan, A.; Campisi, J.; Le Saux, C. J. Secretion of Leukotrienes by Senescent Lung Fibroblasts Promotes Pulmonary Fibrosis. JCI Insight 2019, 4 (24). 10.1172/JCI.INSIGHT.130056.

(14) Pundlik, S. S.; Barik, A.; Venkateshvaran, A.; Sahoo, S. S.; Jaysingh, M. A.; Math, R. G.; Lal, H.; Hashmi, M. A.; Ramanathan, A. Senescent Cells Inhibit Mouse Myoblast Differentiation via the SASP-Lipid 15d-PGJ2 Mediated Modification and Control of HRas. Elife 2024, 13. 10.7554/ELIFE.95229.

(15) Dimri, G. P.; Lee, X.; Basile, G.; Acosta, M.; Scott, G.; Roskelley, C.; Medrano, E. E.; Linskens, M.; Rubelj, I.; Pereira-Smith, O.; Peacocke, M.; Campisi, J. A Biomarker That Identifies Senescent Human Cells in Culture and in Aging Skin in Vivo. Proc. Natl. Acad. Sci. U. S. A. 1995, 92 (20), 9363–9367. 10.1073/PNAS.92.20.9363.

(16) Calió, A.; Zamó, A.; Ponzoni, M.; Zanolin, M. E.; Ferreri, A. J. M.; Pedron, S.; Montagna, L.; Parolini, C.; Fraifeld, V. E.; Wolfson, M.; Yanai, H.; Pizzolo, G.; Doglioni, C.; Vinante, F.; Chilosi, M. Cellular Senescence Markers P16INK4a and P21CIP1/WAF Are Predictors of Hodgkin Lymphoma Outcome. Clin. Cancer Res. 2015, 21 (22), 5164–5172. 10.1158/1078-0432.CCR-15-0508.

(17) Freund, A.; Laberge, R. M.; Demaria, M.; Campisi, J. Lamin B1 Loss Is a Senescence-Associated Biomarker. Mol. Biol. Cell 2012, 23 (11), 2066–2075. 10.1091/MBC.E11-10-0884/ASSET/IMAGES/LARGE/2066FIG4.JPEG.

(18) González-Gualda, E.; Baker, A. G.; Fruk, L.; Muñoz-Espín, D. A Guide to Assessing Cellular Senescence in Vitro and in Vivo. FEBS J. 2021, 288 (1), 56–80. 10.1111/FEBS.15570.

(19) Kuilman, T.; Michaloglou, C.; Mooi, W. J.; Peeper, D. S. The Essence of Senescence. Genes Dev. 2010, 24 (22), 2463–2479. 10.1101/gad.1971610.

(20) Liendl, L.; Grillari, J.; Schosserer, M. Raman Fingerprints as Promising Markers of Cellular Senescence and Aging. GeroScience 2020, 42 (2), 377–387. 10.1007/S11357-019-00053-7.

(21) Ishibashi, S.; Inoko, A.; Oka, Y.; Leproux, P.; Kano, H. Coherent Raman Microscopy Visualizes Ongoing Cellular Senescence through Amide I Peak Shifts Originating from β Sheets in Disordered Nucleolar Proteins. Sci. Rep. 2024, 14 (1). 10.1038/S41598-024-78899-X.

(22) Morgenstern, J.; Fleming, T.; Kadiyska, I.; Brings, S.; Groener, J. B.; Nawroth, P.; Hecker, M.; Brune, M. Sensitive Mass Spectrometric Assay for Determination of 15-Deoxy-Δ12,14- Prostaglandin J2 and Its Application in Human Plasma Samples of Patients with Diabetes. Anal. Bioanal. Chem. 2018, 410 (2), 521–528. 10.1007/s00216-017-0748-1.

(23) Georgiev, D.; Pedersen, S. V.; Xie, R.; Fernández-Galiana, Á.; Stevens, M. M.; Barahona, M. RamanSPy: An Open-Source Python Package for Integrative Raman Spectroscopy Data Analysis. Anal. Chem. 2024, 96 (21), 8492–8500. 10.1021/ACS.ANALCHEM.4C00383/SUPPL_FILE/AC4C00383_SI_003.PDF.

(24) Kumamoto, Y.; Harada, Y.; Takamatsu, T.; Tanaka, H. Label-Free Molecular Imaging and Analysis by Raman Spectroscopy. ACTA Histochem. Cytochem. 2018, 51 (3), 101–110. 10.1267/AHC.18019.

(25) Chandra, A.; Kumar, V.; Garnaik, U. C.; Dada, R.; Qamar, I.; Goel, V. K.; Agarwal, S. Unveiling the Molecular Secrets: A Comprehensive Review of Raman Spectroscopy in Biological Research. ACS Omega 2024, 9 (51), 50049–50063. 10.1021/ACSOMEGA.4C00591.

(26) Di Leonardo, A.; Linke, S. P.; Clarkin, K.; Wahl, G. M. DNA Damage Triggers a Prolonged P53-Dependent G1 Arrest and Long-Term Induction of Cip1 in Normal Human Fibroblasts. Genes Dev. 1994, 8 (21), 2540–2551. 10.1101/gad.8.21.2540.

(27) Hu, X.; Zhang, H. Doxorubicin-Induced Cancer Cell Senescence Shows a Time Delay Effect and Is Inhibited by Epithelial-Mesenchymal Transition (EMT). Med. Sci. Monit. 2019, 25, 3617–3623. 10.12659/MSM.914295.

(28) Robles, S. J.; Adami, G. R. Agents That Cause DNA Double Strand Breaks Lead to P16(INK4a) Enrichment and the Premature Senescence of Normal Fibroblasts. Oncogene 1998, 16 (9), 1113–1123. 10.1038/sj.onc.1201862.

(29) Mosieniak, G.; Sliwinska, M. A.; Alster, O.; Strzeszewska, A.; Sunderland, P.; Piechota, M.; Was, H.; Sikora, E. Polyploidy Formation in Doxorubicin-Treated Cancer Cells Can Favor Escape from Senescence. Neoplasia 2015, 17 (12), 882. 10.1016/J.NEO.2015.11.008.

(30) Srdic-Rajic, T.; Santibañez, J. F.; Kanjer, K.; Tisma-Miletic, N.; Cavic, M.; Galun, D.; Jevric, M.; Kardum, N.; Konic-Ristic, A.; Zoranovic, T. Iscador Qu Inhibits Doxorubicin-Induced Senescence of MCF7 Cells. Sci. Rep. 2017, 7 (1), 1–12. 10.1038/S41598-017-03898-0;TECHMETA.

(31) Campisi, J. Senescent Cells, Tumor Suppression, and Organismal Aging: Good Citizens, Bad Neighbors. Cell 2005, 120 (4), 513–522. 10.1016/j.cell.2005.02.003.

(32) Dilley, T. K.; Bowden, G. T.; Chen, Q. M. Novel Mechanisms of Sublethal Oxidant Toxicity: Induction of Premature Senescence in Human Fibroblasts Confers Tumor Promoter Activity. Exp. Cell Res. 2003, 290 (1), 38–48. 10.1016/S0014-4827(03)00308-2.

(33) Wiley, C. D.; Campisi, J. The Metabolic Roots of Senescence: Mechanisms and Opportunities for Intervention. Nat. Metab. 2021 310 2021, 3 (10), 1290–1301. 10.1038/s42255-021-00483-8.

(34) Olzmann, J. A.; Carvalho, P. Dynamics and Functions of Lipid Droplets. Nat. Rev. Mol. Cell Biol. 2019, 20 (3), 137–155. 10.1038/S41580-018-0085-Z;SUBJMETA.

(35) Liu, K.; Czaja, M. J. Regulation of Lipid Stores and Metabolism by Lipophagy. Cell Death Differ. 2013, 20 (1), 3–11. 10.1038/CDD.2012.63;SUBJMETA.

(36) Vardaki, M. Z.; Gregoriou, V. G.; Chochos, C. L. Biomedical Applications, Perspectives and Tag Design Concepts in the Cell – Silent Raman Window. RSC Chem. Biol. 2024, 5 (4), 273–292. 10.1039/D3CB00217A.

